# Rapid and Specific Identification of Emerging *Trichophyton mentagrophytes* Genotype VII Using an In-House Developed and Validated Real-Time PCR Assay

**DOI:** 10.64898/2026.05.20.726730

**Authors:** Jinjing Zhao, Gabrielle Todd, YanChun Zhu, Sudha Chaturvedi

## Abstract

*Trichophyton mentagrophytes* genotype VII *(Tm*VII) is an emerging sexually transmitted dermatophyte that causes skin infections characterized by inflammatory, erythematous-squamous, painful, and persistent lesions. This genotype is part of the *T. interdigitale/T. mentagrophytes* Species Complex (*TiTm*SC), which comprises 28 genotypes. To enable rapid and specific differentiation of *Tm*VII from other genotypes, a real-time polymerase chain reaction (rt-PCR) assay was developed targeting three unique single-nucleotide polymorphisms in the ITS1 region of *Tm*VII. Assay specificity was further improved by introducing an additional mismatch at the 3’ ends of both forward and reverse primers. The rt-PCR assay demonstrated high sensitivity, with a detection limit of 0.0002 ng of *Tm*VII genomic DNA. The assay was highly specific, with no cross-reactivity observed with either closely or distantly related fungal pathogens when a cycle threshold (Ct) cutoff of 37 was applied. Among 497 mold isolates tested, 47 were confirmed as *Tm*VII by rt-PCR, and the results were fully concordant with conventional ITS-PCR/Sanger sequencing. The rt-PCR assay demonstrated high sensitivity, specificity, reproducibility, and speed, with a turnaround time of one day after DNA extraction, compared with seven to ten days for Sanger sequencing. The first rapid molecular assay developed using TaqMan chemistry for *Tm*VII identification is expected to enhance patient care and support infection control measures.

## INTRODUCTION

*Trichophyton mentagrophytes* genotype VII (*Tm*VII) is an emerging fungal pathogen implicated in tinea corporis and pubogenitalis, a condition that can be transmitted via sexual contact [1]. Tinea corporis is a common superficial skin infection that typically affects the trunk and extremities and is generally mild and easily treatable. In contrast, tinea of the pubogenital region is rare and primarily reported in tropical regions. Recently, a distinct form of tinea, characterized by annularly inflamed, scaly, painful, itchy, and persistent skin lesions caused by *Tm*VII, has been identified [2, 3]. A 2015 study documented three severe genital region infections, each with a history of sexual contact in Southeast Asia prior to symptom onset, attributed to a previously unknown genotype of *T. mentagrophytes* likely corresponding to *Tm*VII [4]. In 2019, a German research group identified a new genotype of *T. mentagrophytes* from 37 cases of sexually transmitted infection, classifying it as *Tm*VII [5] with subsequent reports of other genotypes [6]. This pathogen has since spread globally to France [1], Italy [7], the United Kingdom [8], Spain [9], China [10] and the United States [2]. There is an urgent need for a simple and rapid method to identify *Tm*VII to enable appropriate treatment and prevent further spread.

*Tm*VII is a member of the *T. interdigitale/T. mentagrophytes* Species Complex (*TiTm*SC). Genotype-level identification is essential because different members of the *TiTm*SC exhibit distinct mechanisms of transmission, preferred infection sites, and drug resistance profiles [11]. The standard method for *Tm*VII identification involves conventional polymerase chain reaction (PCR) targeting the internal transcribed spacer (ITS) region, followed by Sanger sequencing and BLAST analysis. This process is both time-consuming and costly. Alternative detection methods have been developed to reduce the resources required for dermatophyte identification, including loop-mediated isothermal amplification (LAMP), rt-quantitative PCR (qPCR) [12], and DermaGenius 2.0, a commercially available, complete multiplex rt-PCR test introduced in 2018. [13]. However, these methods identify *T. mentagrophytes* to the species level but cannot differentiate among genotypes within the *TiTm*SC.

To address these limitations, in the current study, we developed a highly sensitive, *Tm*VII-specific real-time PCR (rt-PCR) assay based on unique *Tm*VII single nucleotide polymorphisms (SNPs) identified among the ITS sequences from all 28 known genotypes of *TiTm*SC [14]. The *Tm*VII rt-PCR assay was validated for sensitivity, specificity and reproducibility using gDNA from *Tm*VII, closely related *TiTm*SC members, and distantly related dermatophytes, molds and yeasts. Prospective comparisons of *Tm*VII rt-PCR with ITS-PCR/sequencing demonstrated that accurate *Tm*VII identification could be achieved within one day of DNA extraction, compared to 7 to 10 days required for identification by the ITS-PCR/sequencing assay. This assay substantially reduces turnaround time, which is expected to improve patient care and facilitate infection control measures.

## MATERIALS AND METHODS

### Strains and DNA

All *Tm*VII clinical isolates used in this investigation were collected from multiple health care facilities in New York City and surrounding counties between 2023 and 2025. Closely related isolates in the *TiTm*SC, including *T. interdigitale* I, *T. interdigitale* II, *T. interdigitale* XI, *T. mentagrophytes* II* (*Tm*II*), *T. mentagrophytes* III* (*Tm*III*), *T. mentagrophytes* IV (*Tm*IV), *T. mentagrophytes* XVIII (*Tm*XVIII), and the new species *T. indotineae* (formerly *T. mentagrophytes* VIII), as well as other non-dermatophyte molds used in this study were identified by our in-house ITS-PCR/Sanger sequencing protocol. Yeast isolates were identified in-house by MALDI-TOF MS.

DNA extracted from dermatophytes, other molds and yeasts, as described previously [11], was stored at -80°C until use in the *Tm*VII rt-PCR assay development and validation.

### Primers, probe design and real-time PCR assay set up

A multiple alignment of the 28 genotypes of *TiTm*SC [14] revealed three unique single-nucleotide polymorphisms (SNPs) specific to *Tm*VII within the ITS1 region (Supplementary Fig. 1A). Primers were designed such that two of the SNPs unique to *Tm*VII were located at the 3’ ends of the forward (C>G) and reverse (C>T) primers. The rt-PCR probe was designed to bind the third unique SNP (C>T). To further enhance primer specificity for *Tm*VII, an additional mismatch was introduced at the 3’ region of the forward (‘T’ highlighted in green) and reverse primers (‘A’ highlighted in red) (Supplementary Fig. 1A) as described previously [15]. Primer specificity was evaluated using conventional PCR with 0.2 ng genomic DNA (gDNA) per PCR reaction from three isolates of *Tm*VII, and one isolate each of *T. interdigitale* I and II, *Tm*III*, *Tm*VI, and *T. indotineae*, with the following cycling condition: 95°C for 60 s, 45 cycles of 95°C for 15 s, 62°C for 15 s and 68°C for 60 s, and then final extension for another 5 mins. The primers generated a 113-bp amplicon exclusively with gDNA from *Tm*VII (Supplementary Fig. 1B). *Tm*VII rt-PCR reactions contained 1X PerfeCTa Multiplex qPCR ToughMix (Quanta Biosciences, Part # 95147-05K), 0.4 µM each of forward and reverse primer, 0.1 µM of probe and 2 µL of isolate DNA ranging from 0.0001 ng to 10 ng/µL in 20 µL reaction volume. Reactions were run in an optical 96-well reaction plate (Applied Biosystems, Cat. No. 4481192) on a QuantStudio 7 Pro Dx (Applied Biosystems) with the following cycling conditions: 95°C for 60 s, 45 cycles of 95°C for 15 s, and 62°C for 50 s. Cycle threshold (Ct) values were observed only from *Tm*VII and not for any other dermatophytes tested (Supplementary Fig. 1C). These preliminary results were the basis for ordering the bulk of primers and probe from Integrated DNA Technologies, Inc. (Coralville, Iowa). Sequences for primers and probe were as follows: V2670 (*Tm*VII_F1), 5’-CAGGCCGGAGGCTGGTCG-3’; V2671 (*Tm*VII_R2), 5’-GCCCGCCGAGGCAACCTAA-3’; V2672 (*Tm*VII_P), 5’-/56 FAM/TGTGCGCCGGCCGTACCGTCC/3IABkFQ/ -3’.

### Analytical sensitivity, specificity, reproducibility, and accuracy

To evaluate the *Tm*VII rt-PCR sensitivity, gDNA of one isolate of *Tm*VII was serially diluted and tested in triplicate in the rt-PCR assay on three different days. Linear regression analysis was performed across six logs of gDNA concentrations. The slope (*s*), correlation coefficient (*R*^*2*^), and the amplification efficiency (*e*) of the PCR assay were calculated using an equation as provided [16]. Assay specificity was determined by analyzing gDNA from 72 fungal pathogens both closely and distantly related to *Tm*VII. Assay reproducibility was performed using low, medium, and high gDNA concentrations of three isolates of *Tm*VII in triplicate on the same and over three different days of testing. Assay verification was performed by testing a blinded panel consisting of 31 *Tm*VII isolates at various gDNA concentrations, with one concentration near the limit of detection (LOD) and 17 others at higher DNA concentrations.

### Assay Performance

The assay performance was evaluated using both retrospective and prospective fungal pathogens received in the laboratory. The retrospective studies included gDNA from 38 *Tm*VII isolates received in the laboratory since its emergence in 2023, and 140 other genotypes of *TiTmSC* isolates received from 2017 to 2025, and 53 isolates of non-dermatophytes species. The extracted gDNA was diluted 1 to 100-fold without measuring DNA concentrations, and 2 µL of diluted DNA was directly used in rt-PCR reaction. To determine turnaround time, a total of 266 mold isolates received in the laboratory for fungal identification by ITS PCR/Sequencing and BLAST search were also subjected to *Tm*VII rt-PCR assay prospectively using the same method described above for the retrospective study.

### Statistical analysis

Graph Pad Prism10.4.1 for Windows was used for statistical analysis. The mean, standard deviation, and percent coefficient of variance (CV) for Ct value were calculated for assay sensitivity and reproducibility. A Student’s *t* test was used for analysis of the means, and a *P* value ≤ 0.05 was considered statistically significant.

## Results

### *Tm*VII rt-PCR is Highly Sensitive, Specific, and Reproducible

The *Tm*VII rt-PCR assay demonstrated high sensitivity, with a limit of detection (LOD) of 0.0002 ng gDNA per PCR reaction and a Ct cutoff value of 37 using 45 PCR cycles (Supplementary Table 1). The assay exhibited linearity across six orders of magnitude. The slope (*s*), correlation coefficient (*R*^2^), and amplification efficiency (*e*) were excellent (Fig. 1). The assay was highly reproducible with consistent *Ct* values obtained for gDNA concentrations tested from three different *Tm*VII isolates within the same day and on three different days of testing (Supplementary Tables 2 A & 2 B). The assay was highly specific, with no cross-reactivity observed to closely or distantly related fungal pathogens when a Ct cutoff value of 37 was set (Supplementary Table 3). Ct values exceeding 37 were interpreted as negative for *Tm*VII. From a panel of blinded samples, the assay readily detected *Tm*VII at DNA concentrations ranging from 0.002 to 20 ng per PCR reaction. A few dermatophytes other than *Tm*VII yielded low amplicons with Ct values beyond 37 at a very high concentration of gDNA (20 ng per PCR reaction). The rest of the molds and yeasts produced negative results, confirming the high accuracy of this *Tm*VII rt-PCR (Supplementary Table 4).

**Table 1.** *Tm*VII rt-PCR assay performance and turnaround time.

| rt-PCR<br>(days to<br>ID = 1) | ITS-PCR/Sequencing<br>(days to ID = 7-10) |  | Accuracy | Sensitivity | Specificity | PPV (%) | NPV (%) |
| --- | --- | --- | --- | --- | --- | --- | --- |
|  | Positive | Negative |  |  |  |  |  |
| Positive | 9 (TP) | 0 (FP) | 100% | 100% | 100% | 100% | 100% |
| Negative | 0 (FN) | 257 (TN) |  |  |  |  |  |

**Figure 1.**
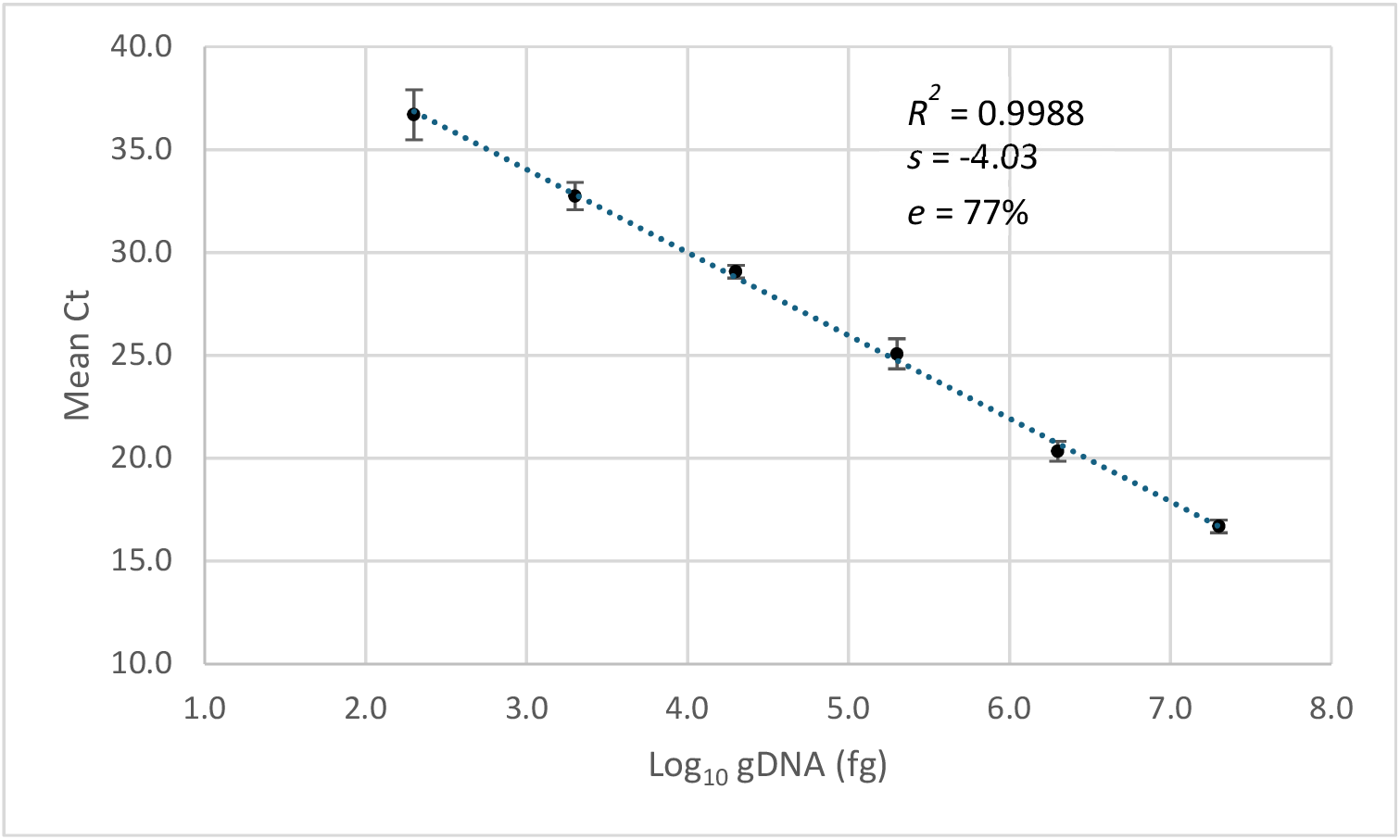
*Tm*VII rt-PCR assay sensitivity. A linear regression analysis was performed to determine assay sensitivity. The gDNA from one isolate of *Tm*VII was serially diluted, and mean *Ct* values from duplicate runs on three different days for each dilution were plotted. The assay exhibited linearity across six orders of magnitude. The slope (*s*), correlation coefficient (*R*^2^), and amplification efficiency (*e*) are shown.

### *Tm*VII rt-PCR Revealed High Performance

After establishing the assay, its performance was assessed by retrospectively testing gDNA from 231 fungal pathogens and comparing the rt-PCR results with the organism ID obtained by Sanger sequencing of the ITS region. Each gDNA sample was diluted 1:100, and 2 µL was analyzed in each *Tm*VII rt-PCR assay. Of the 231 fungal pathogens, 38 were positive for *Tm*VII with Ct values between 17 and 22. A limited number of other dermatophytes within the *TiTm*SC group also produced Ct values, but these Ct values ranged from 39 to 45 which exceeds our established cutoff of 37. No other molds or yeasts were detected by the *Tm*VII rt-PCR assay (Fig. 2). These findings further validate our Ct cutoff value and demonstrate that this rt-PCR assay is highly accurate for *Tm*VII identification (Supplementary Table 5).

**Figure 2.**
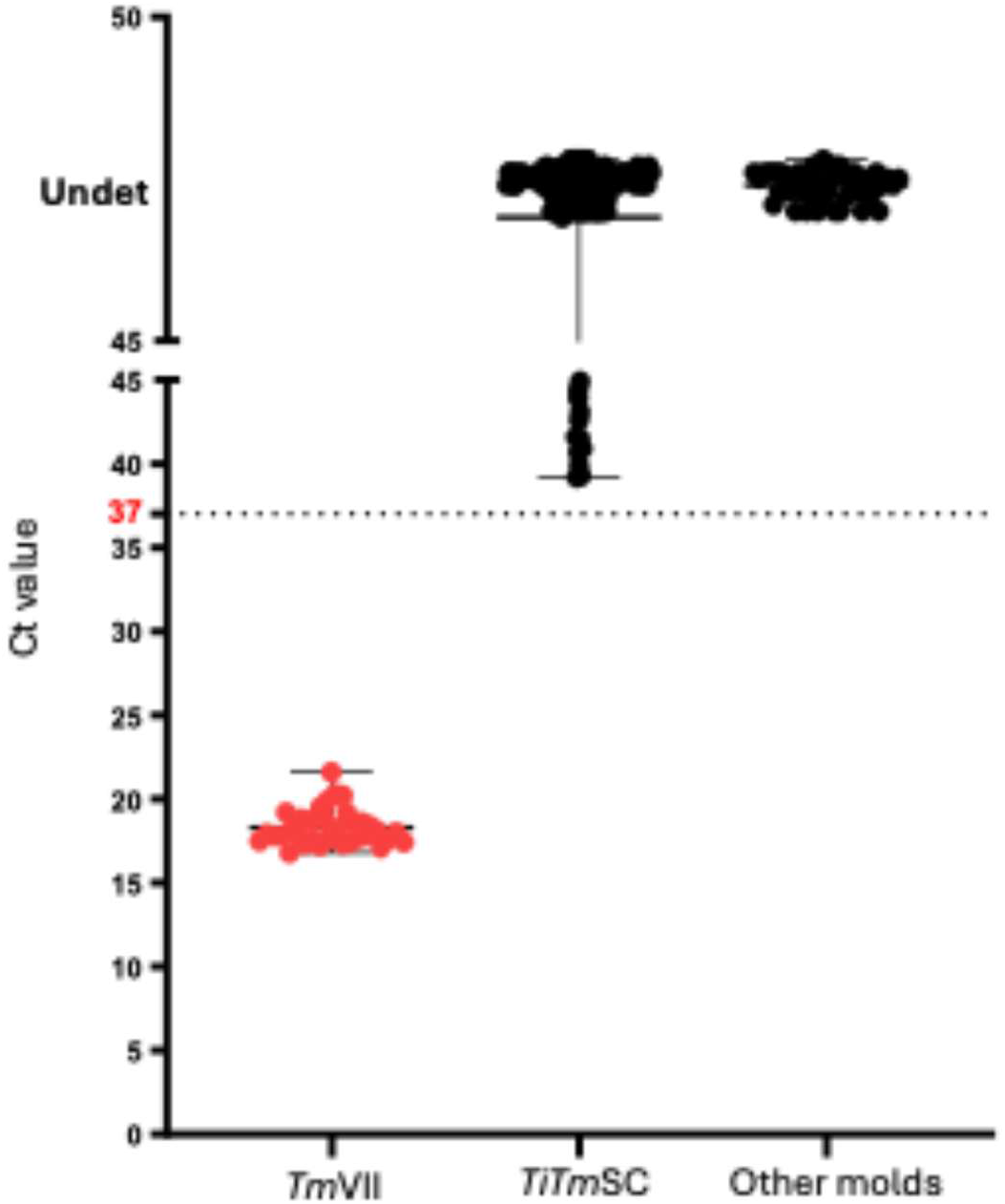
*Tm*VII rt-PCR assay performance using retrospective mold DNA samples. The gDNA from 231 molds was diluted 100-fold with water and 2 µL of the diluted DNA was run in the rt-PCR assay in duplicate. All *Tm*VII isolates yielded Ct values below the cutoff value of 37, while a few species within the *TiTm*SC yielded Ct values above the cutoff, and other molds yielded no Ct, confirming the high degree of accuracy of the *Tm*VII rt-PCR assay.

The turnaround time of the *Tm*VII rt-PCR assay was compared to Sanger sequencing using prospective samples. A total of 266 mold isolates received between July 1, 2025, and September 16, 2025, were processed for DNA extraction and diluted 1:100 as previously described. Two microliters of diluted DNA were used for the *Tm*VII rt-PCR assay. Of the 266 mold isolates, nine were identified as *Tm*VII by rt-PCR within one day of DNA extraction, with Ct values ranging from 16 to 29. Eight isolates produced Ct values between 40 and 45; among these, five were confirmed as *T. indotineae*, two as *T. interdigitale* II, and one each as *T. mentagrophytes* II* and *Cystobasidium slooffiae* (Supplementary Table 6) by Sanger sequencing. The *Tm*VII rt-PCR assay did not cross-react with other molds. *Tm*VII identification was achieved within one day of gDNA extraction using rt-PCR, whereas Sanger sequencing took 7 to 10 days for identification (Table 1). Thus, this rt-PCR assay is highly accurate, specific, and allows for rapid identification of *Tm*VII isolates.

## Discussion

*Trichophyton mentagrophytes* genotype VII (*Tm*VII) is an emerging dermatophyte associated with sexual transmission among men who have sex with men [1, 17]. Since its identification, *Tm*VII has spread globally, prompting concern among healthcare professionals. The first case of *Tm*VII infection in the United States was described in July 2024 [2], with several additional cases subsequently reported [3], and recently *Tm*VII cases in New York have rapidly increased with possible clonal expansion of this dermatophyte [11]. Prompt recognition and appropriate treatment of *Tm*VII infections, typically with oral terbinafine, are essential for improving patient outcomes. *Tm*VII infections are frequently misdiagnosed as non-infectious skin conditions, such as psoriasis, or as other sexually transmitted infections [2]. Delayed treatment may result in secondary bacterial infection and ongoing transmission [1]. Consequently, rapid and accurate identification of *Tm*VII is vital for effective patient management. To address this need, a TaqMan chemistry, probe-based, real-time PCR (rt-PCR) assay was developed for the rapid identification of suspected *Tm*VII isolates received in the laboratory. This assay demonstrated high specificity, speed, and accuracy in the identification of *Tm*VII.

The genotypes within the *TiTm*SC are generally closely related, limiting the availability of single-nucleotide polymorphisms (SNPs) that can distinguish genotypes within the complex. This presents challenges for designing an assay that is both specific and sensitive. To improve sensitivity, we chose the ITS region, a gene present in multiple copies. For greater specificity, we focused on the ITS1 region and SNPs unique to *Tm*VII when designing primers and probes. We also increased specificity by adding an extra mismatch at the 3’ ends of both primers, making it harder for other *TiTm*SC genotypes to initiate amplification, as described before for *T. indotineae* identification [15]. This approach, called “Polymerase Extension Block,” prevents the polymerase from starting polymerization if there are mismatches at the last 1-5 nucleotides of the primer’s 3’ end. This acts as a barrier and improves the assay’s specificity for *Tm*VII. The assay gave high amplicon yields for *Tm*VII with as little as 0.0002 ng of gDNA per PCR reaction, and a cycle threshold (Ct) cutoff of 37. In comparison, only low amplicon yields were seen for a few *TiTm*SC species, even when using 20 ng of gDNA per PCR reaction, and their Ct values were above 37. Out of 497 molds tested by rt-PCR, 47 were confirmed as *Tm*VII, matching Sanger sequencing results for the ITS region. A few isolates with high Ct values (over 37) were identified as other *TiTm*SC genotypes by Sanger sequencing, showing that the rt-PCR assay is highly specific without losing sensitivity.

Given the rapid spread of *Tm*VII in New York and globally, the developed rt-PCR assay enables differentiation of *Tm*VII from other genotypes within *TiTm*SC and from *T. indotineae*, thereby facilitating timely initiation of appropriate therapy. Current literature indicates that *Tm*VII infection responds to the first-line drug terbinafine, whereas *T. indotineae* does not [18, 19]. In vitro susceptibility data further indicate that *Tm*VII is susceptible to all tested antifungal agents, including terbinafine [11]. The *Tm*VII rt-PCR assay for isolates yields results within one day of DNA extraction, compared to 7–10 days required for conventional ITS PCR and Sanger sequencing. This highly sensitive, specific, and rapid assay is anticipated to advance diagnostic capabilities, improve patient outcomes, and strengthen infection control measures. Future studies are planned with primary skin samples, in collaboration with healthcare providers, which will further enhance the diagnostic process and patient care.

## Figure Legends

**Supplementary Figure 1.**
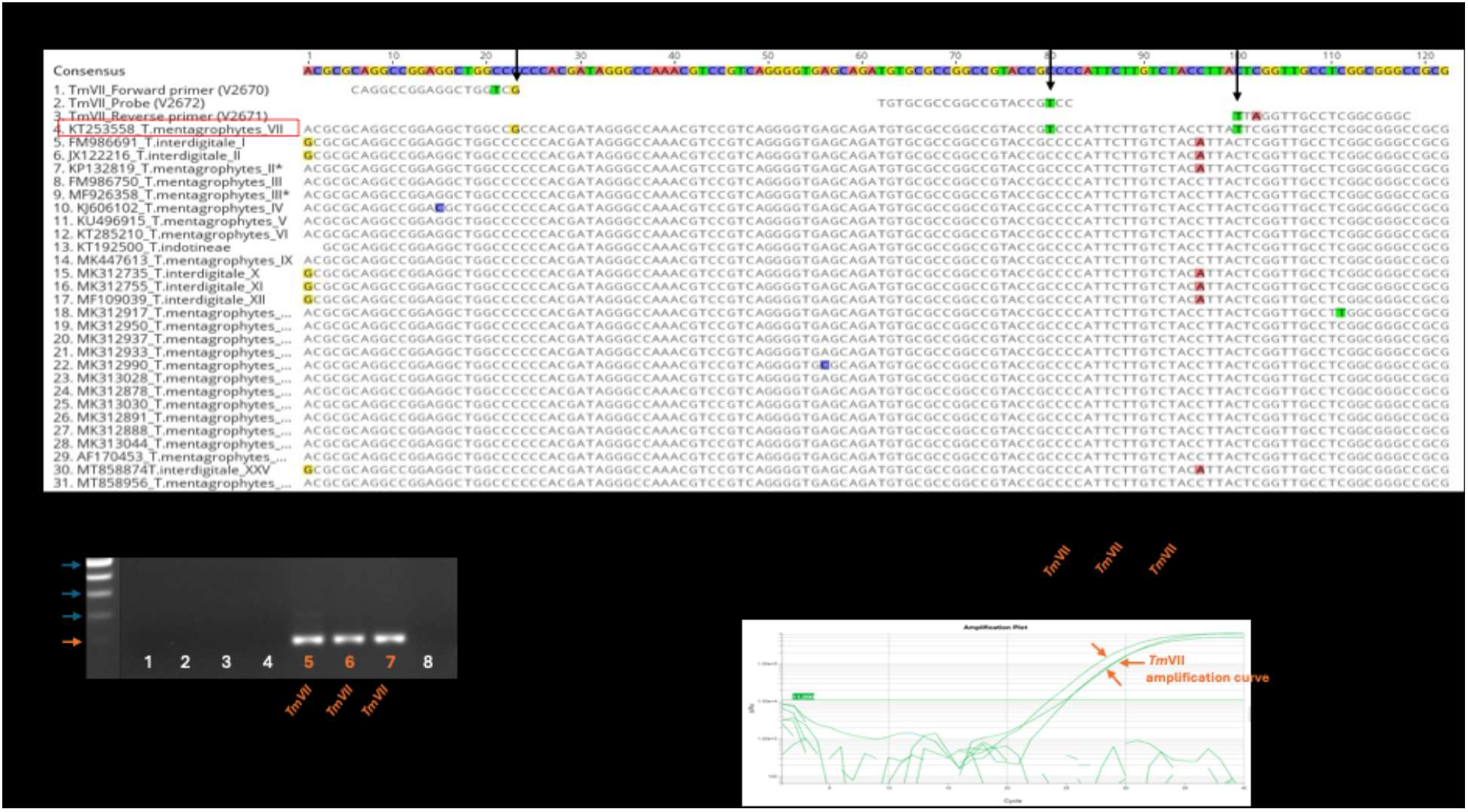
**A)** Multiple alignment of the ITS region of all 28 genotypes in the *TiTm*SC revealed three unique SNPs (black arrows) in *Tm*VII, which were used for primer and probe design. Additionally, a mismatch was created at the 3’ end of the forward (T in green) and reverse (A in red) primers to increase assay specificity. **B)** Conventional PCR with the forward and reverse primers produced a 113-bp amplicon for *Tm*VII, but not for other closely related dermatophytes, confirming primer specificity. **C)** Real-time PCR yielded Ct values only for *Tm*VII, and not for other dermatophytes, confirming primer and probe specificity.

## Authors Contributions

JJZ designed and performed rt-PCR assays, analyzed data, prepared graphs and tables, and wrote the first draft of the manuscript. YCZ performed ITS PCR, analyzed Sanger sequences and identified *Tm*VII and other molds. GT helped blind the panel, provided ITS sequences of 28 genotypes of *TiTm*SC, and edited the manuscript. SC conceived the study and extensively edited the manuscript. All authors contributed to the manuscript and approved the submitted version.

## Acknowledgements

We thank Wadsworth Center Media and Tissue Culture Core and Advanced Genomic Technologies Core for media and DNA sequencing. We also thank Dr. Lisa A. Biega for critical reading of the manuscript. This study was partially supported by the Centers for Disease Control and Prevention Epidemiology and Laboratory Capacity Antibiotic Resistance Laboratory Network Grant NU51CK000372.

## References

1. Jabet A, Dellière S, Seang S, Chermak A, Schneider L, Chiarabini T, Teboul A, Hickman G, Bozonnat A, Brin C et al: Sexually Transmitted Trichophyton mentagrophytes Genotype VII Infection among Men Who Have Sex with Men. Emerg Infect Dis 2023, 29(7):1411–1414.

2. Caplan AS, Sikora M, Strome A, Akoh CC, Otto C, Chaturvedi S, Zampella JG: Potential Sexual Transmission of Tinea Pubogenitalis From TMVII. JAMA Dermatol 2024, 160(7):783–785.

3. Zucker J CA, Gunaratne SH, et al.: Notes from the Field: Trichophyton mentagrophytes Genotype VII — New York City. Morbidity and Mortality Weekly Report 2024, 73:985–988.

4. Luchsinger I, Bosshard PP, Kasper RS, Reinhardt D, Lautenschlager S: Tinea genitalis: a new entity of sexually transmitted infection? Case series and review of the literature. Sex Transm Infect 2015, 91(7):493–496.

5. Kupsch C, Czaika VA, Deutsch C, Gräser Y: Trichophyton mentagrophytes - a new genotype of zoophilic dermatophyte causes sexually transmitted infections. Journal Der Deutschen Dermatologischen Gesellschaft 2019, 17(5):493–502.

6. Heidemann S, Monod M, Graser Y: Signature polymorphisms in the internal transcribed spacer region relevant for the differentiation of zoophilic and anthropophilic strains of Trichophyton interdigitale and other species of T. mentagrophytes sensu lato. Br J Dermatol 2010, 162(2):282–295.

7. Rossi L, Sorrentino A, Signoretto PC, Gaibani P: Genome characterization of Trichophyton mentagrophytes genotype VII strain PG12DES from Italy. Med Mycol 2025, 63(6).

8. Tan XL, Lambourne J, Borman AM, Abdolrasouli A, Goldsmith P, Cunningham M, Gkini MA: Trichophyton mentagrophytes genotype VII sexually transmitted infection in the UK: where are we now? Br J Dermatol 2025, 193(3):566–568.

9. Descalzo V, Martin MT, Alvarez-Lopez P, Garcia-Perez JN, Alcazar-Fuoli L, Lopez-Perez L, Tellez-Velasco D, Carrillo A, Sulleiro E, Falco V et al: Trichophyton mentagrophytes Genotype VII and Sexually Transmitted Tinea: An Observational Study in Spain. Mycoses 2025, 68(4):e70049.

10. Zhang Y, Xie W, Liu W, Li X, Liang G: Epidemiological and Clinical Profile Analysis of Trichophyton mentagrophytes ITS Genotype VII Infected Dermatomycosis: An Emerging Sexually Transmitted Pathogen. Mycoses 2025, 68(6):e70075.

11. Todd Gabrielle C, Vaida V, O’Brien B, Chaturvedi S: The emergence of superficial dermatophytosis due to Trichophyton indotineae and Trichophyton mentagrophytes genotypes VII and II* in New York: a need for comprehensive testing approaches. Journal of Clinical Microbiology 2026, 0(0):e00156–00126.

12. Müs Tak HK, Ünal G, Müs Tak I: Detection of Microsporum canis and Trichophyton mentagrophytes by loop-mediated isothermal amplification (LAMP) and real-time quantitative PCR (qPCR) methods. Vet Dermatol 2022, 33(6):516–522.

13. Aho-Laukkanen E, Mäki-Koivisto V, Torvikoski J, Sinikumpu SP, Huilaja L, Junttila IS: PCR enables rapid detection of dermatophytes in practice. Microbiol Spectr 2024, 12(11):e0104924.

14. Normand AC, Moreno-Sabater A, Jabet A, Hamane S, Cremer G, Foulet F, Blaize M, Delliere S, Bonnal C, Imbert S et al: MALDI-TOF Mass Spectrometry Online Identification of Trichophyton indotineae Using the MSI-2 Application. J Fungi (Basel) 2022, 8(10).

15. Baron A, Hamane S, Gits-Muselli M, Legendre L, Benderdouche M, Mingui A, Ghelfenstein-Ferreira T, Alanio A, Dellière S: Dual quantitative PCR assays for the rapid detection of Trichophyton indotineae from clinical samples. Med Mycol 2024, 62(7).

16. Ginzinger DG: Gene quantification using real-time quantitative PCR: an emerging technology hits the mainstream. Exp Hematol 2002, 30(6):503–512.

17. Jabet A, Favier M, Normand AC, Pourcher V, Muller VL, Monsel G: [Trichophyton mentagrophytes ITS genotype VII: transmission in the absence of visible lesions?]. Dermatologie (Heidelb) 2025, 76(10):640–643.

18. Caplan AS, Todd GC, Zhu Y, Sikora M, Akoh CC, Jakus J, Lipner SR, Graber KB, Acker KP, Morales AE et al: Clinical Course, Antifungal Susceptibility, and Genomic Sequencing of Trichophyton indotineae. JAMA Dermatol 2024, 160(7):701–709.

19. Gold JAW, Lipner SR: The Rise of Antifungal-Resistant Dermatophyte Infections: What Dermatologists Need to Know. Cuts 2025, 115(5):151–154.

